# Expanded utility of the R package qgg with applications within genomic medicine

**DOI:** 10.1101/2022.09.03.506466

**Authors:** Palle Duun Rohde, Izel Fourie Sørensen, Peter Sørensen

## Abstract

**Summary:** Here, we present an expanded utility of the R package qgg for quantitative genetic and genomic analyses of complex traits and diseases. One of the major updates of the package is, that it now includes five different Bayesian Linear Regression (BLR) models, which provide a unified framework for mapping of genetic variants, estimation of heritability and genomic prediction from either individual level data or from genome-wide association study (GWAS) summary statistics. To showcase some of the novel implementations, we analysed two quantitative trait phenotypes, body mass index and standing height from United Kingdom Biobank (UKB). We compared genomic prediction accuracies from single and multiple trait models, showed accurate estimation of genomic parameters, illustrate how a BLR model can be used to fine map potential causal loci, and finally, provide an extension of gene set enrichment analyses based on the BLR framework. With this release, the qgg package now provides a wealth of the commonly used methods in analysis of complex traits and diseases, without the need to switch between software tools and data formats.

**Availability:** Our methodology is implemented in the publicly available R software package qgg using fast and memory efficient algorithms in C++ and is available from CRAN or as a developer version at our GitHub page (https://github.com/psoerensen/qgg). Notes on the implemented statistical genetic models, tutorials and example scripts are available from our accompanied homepage https://qganalytics.com/.

**Supplementary information:** Supplementary data are available online.

## 1 Introduction

With the increasing availability of large genetic biobanks and as each biobank increase in number of collected samples, there is a demand for software that allow for robust and efficient genomic analyses. Tools as PLINK (Chang *et al.*, 2015), GCTA (Yang, Lee, *et al.*, 2011), LDpred (Privé *et al.*, 2021; Vilhjálmsson *et al.*, 2015), LDAK (Zhang *et al.*, 2021; Speed *et al.*, 2017; Speed and Balding, 2014) and PRSice (Euesden *et al.*, 2015) have changed the way researchers around the globe conduct genetic analyses of human complex traits and diseases. Here, we present a new release of the R package qgg (Rohde *et al.*, 2020), which now has been expanded to include a range of commonly used methods such as LD Score Regression (LDSC), best linear unbiased prediction (BLUP) estimation from ordinary least squares (OLS), a range of different Bayesian shrinkage models for gene mapping and trait prediction, multiple-trait genetic risk profiling and much more. With a user-friendly interface, qgg offers a unified tool for quantitative genetic analysis of complex traits and diseases. Accompanied with the second major release of qgg follows a repository (www.qganalytics.com) containing software user guides and detailed notes about the statistical genetic models and methods implemented in qgg.

Human complex traits and disease varies greatly in how their genetic architecture is shaped, e.g., in terms of effect size distribution, number of causal variants, non-linear genetic effects, etc (Timpson *et al.*, 2018). As every statistical tool has its underlying statistical assumptions which necessarily reflect the type of genetic architecture that can be detected, it can be useful to exploit different statistical algorithms e.g., for gene mapping or genomic risk prediction. Bayesian linear regression (BLR) models have been proposed as a unified framework for mapping of genetic variants, estimation of heritability and genomic prediction (Moser *et al.*, 2015; Lloyd-Jones *et al.*, 2019; Vilhjálmsson *et al.*, 2015; Ehsani *et al.*, 2012, 2016; Sørensen *et al.*, 2015). BLR models account for the underlying genetic architecture by allowing the true genetic signal to be heterogeneous distributed over the genome (i.e., some regions have stronger genetic signal than others). Because BLR models fit all genetic markers simultaneously and account for linkage disequilibrium (LD) between markers, they often have greater power to detect causal associations, and find less false negatives (Lloyd-Jones *et al.*, 2019). The gain in statistical power to detect causal variants subsequently increase the accuracy of genomic prediction. In this release of the qgg package, a range of different BLR models has been implemented.

To showcase some of the new features within qgg we analysed two quantitative trait phenotypes, namely body mass index (BMI) and standing height, using data from United Kingdom Biobank (UKB) (Bycroft *et al.*, 2018). Many quantitative traits and multifactorial diseases are genetically correlated (Bulik-Sullivan *et al.*, 2015), hence, we and others, have previously shown that the accuracy of genetic prediction can be improved by leveraging the genetic correlations among such phenotypes (Maier *et al.*, 2018; Rohde *et al.*, 2021). Here, we report additional improvements in prediction accuracy by combining the theory from multiple-trait models with the BLR models, by jointly analysing standing height and BMI with males and females treated as individual phenotypes (i.e., four traits in total). We further show an example of genetic fine mapping of putative causal variants using a BLR model, and how we can partition genomic parameters by utilising a multiple trait BLR model. Finally, we extend the traditionally gene set enrichment analyses by integrating the BLR framework.

## 2 Overview of qgg

The R package qgg provides a user-friendly and streamlined infrastructure (Figure 1) for genetic analysis.

**Figure 1.**
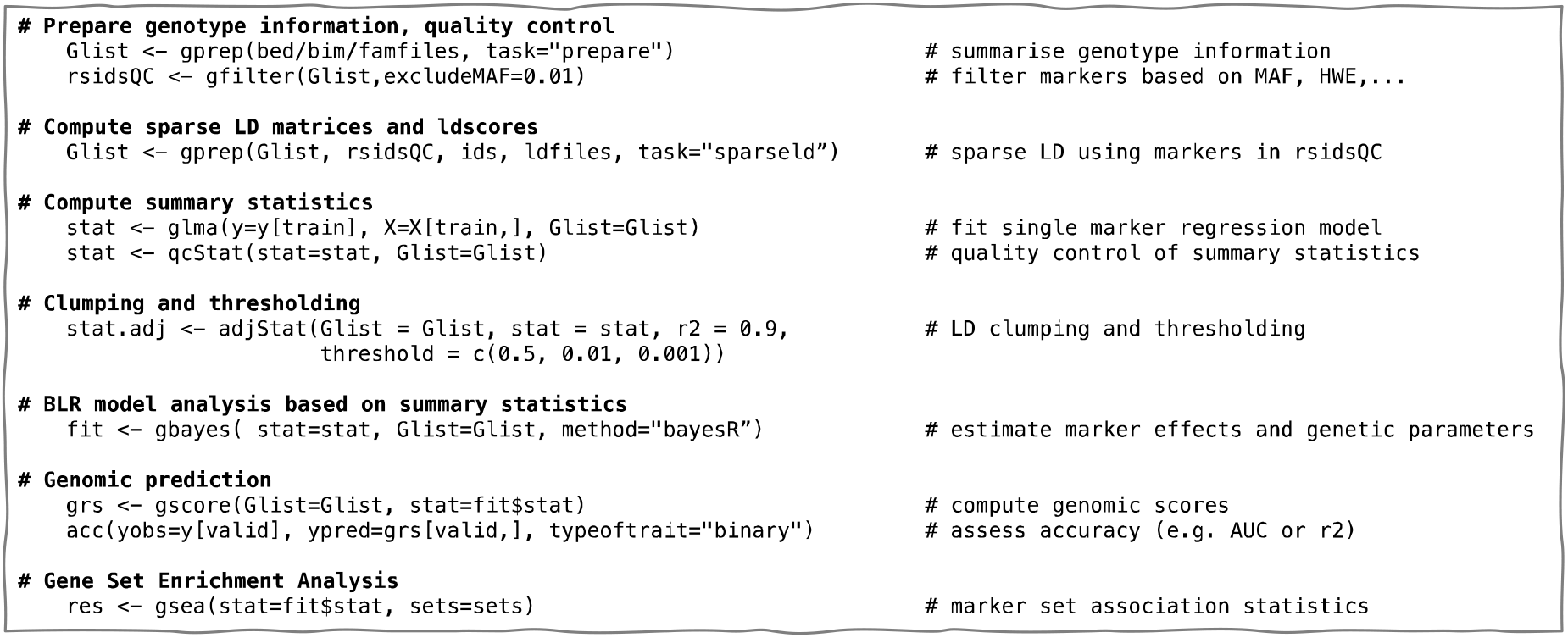
Simple and streamlined workflow for genetic analysis of complex trait that takes PLINK files as input. The output from each function is designed to match the input format of downstream analyses.

Initially, PLINK files are processed to construct a Glist-object that summarises information about the genetic data and computes genotype and allele frequencies, missingness etc. This information can be used to perform standard quality control of the genetic data (Marees *et al.*, 2018), and to compute sparse LD matrices and LD scores. After the initial pre-processing, GWAS summary statistics can directly be computed, which can be adjusted either using clumping and thresholding (C+T) or one of the implemented BLR models. Finally, the GWAS summary statistics can be used to construct genomic scores (GS) or to identify biologically relevant gene sets enriched for associated variants.

## 3 Materials and methods

### 3.1 Genetic and phenotypic data

Genetic and phenotypic data were obtained from the United Kingdom Biobank (UKB), in which data has been collected for more than 500.000 individuals aged 37-73 years (Bycroft *et al.*, 2018). Genotyping details has been described previously (Bycroft *et al.*, 2018). To obtain a genetic homogeneous study population we restricted our analyses to unrelated British, Caucasians and excluded individuals with more than 5,000 missing genotypes or individuals with autosomal aneuploidy (*n*=335,744). For the purpose of presenting the novel utilities of the qgg-package we used the genotyped variants and excluded those with minor allele frequency <0.01, deviation from Hardy-Weinberg equilibrium (*P*-value <1 · 10^-12^), if genetic variants were located within the major histocompatibility complex, had allele ambiguous (*i.e.,* GC or AT), was multi-allelic or an indel (Marees *et al.*, 2018). This resulted in a total of 533,679 genotyped variants.

Standing height (data field 50) and body mass index (BMI, data field 21001) was used as example traits. Prior to analyses the two traits were for each sex separately adjusted for age, UKB assessment centre and the first ten genetic principal components following inverse rank normalisation.

Linkage disequilibrium (LD) among quality-controlled genotype variants was computed using a random selection of 50,000 UKB participants in window sizes of 2000 variants on each side for a genotyped marker. LD was computed as the observed correlation among genotyped variants as 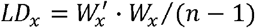, where *x* was the index of the set of variants of which LD was computed, *W* was a centred and scaled (to mean zero and variance one) genotype matrix, and *n* was the number of individuals.

### 3.2 Genomic scores (GS)

A genomic score (GS) captures an individual’s genetic predisposition for a given multifactorial trait, and can be computed as in Eq. 1 (Dudbridge, 2013; Purcell *et al.*, 2009):

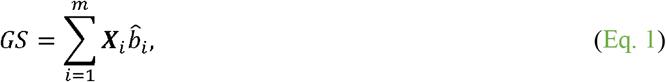

where ***X**_i_* denotes the *i*-th genotype (encoded as 0, 1, 2 counting the number of the alternative allele), and 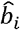 is the marker effect on the corresponding phenotype (for binary outcomes then 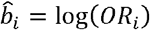, where OR is the odds ratio of the *i*-th genetic variant). The marker effects 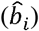 are typically marginal effects from linear/logistic regressions (Chang *et al.*, 2015), linear mixed models (Yu *et al.*, 2006; Zhou and Stephens, 2012), or using Bayesian Linear Regressions (BLR) (Loh *et al.*, 2015; Mbatchou *et al.*, 2021). In absence of shrinkage of the marginal effects, several different approaches can be applied to select genetic variants used in constructing GS, which is described below.

#### 3.2.1 Clumping and thresholding

GS are commonly computed using clumping and thresholding (C+T) (Euesden *et al.*, 2015), that involves sorting all genetic variants based on their association *P*-value statistics and removing those variants that are strongly correlated with variants with *P*-values below a certain threshold (Choi *et al.*, 2020).

Important parameters to optimize for C+T is the squared correlation among genetic variants (*r*^2^) and the *P*-value significance level (*P*) (Privé *et al.*, 2019). Then, for different values of *r*^2^ and *P* the C+T predictors are computed using Eq. 1.

#### 3.2.2 Bayesian linear regression models for marker effects shrinkage

Complex traits and multifactorial diseases are likely highly polygenic, with hundreds to thousands of causal variants, most frequently with small effect sizes (Manolio *et al.*, 2009; Timpson *et al.*, 2018). BLR models have become increasingly popular, as this collection of models jointly (re)-estimate marker effects of millions of variants sampling marker effects from prior known distributions while simultaneously accounting for the correlation structure among genetic markers (i.e., LD) (Moser *et al.*, 2015).

Different prior distributions of marker effects have been proposed (Gianola, 2013; Habier et al., 2011), and can be explained as a two-stage distribution, in which each marker has its own prior:

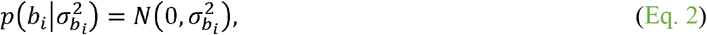

and a prior distribution of the variance *per se*:

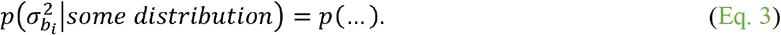

A range of different BLR models assuming different prior distributions has been implemented in the qgg package, which shortly are described below.

##### Prior marker variance BayesN

The simplest marker variance prior, is to assume that all marker effects have the same Gaussian variance (e.g., Meuwissen et al. (2001)). Assigning a Gaussian prior to *b* implies that the posterior means are equivalent to the BLUP estimates:

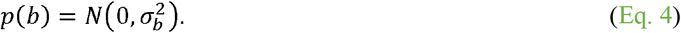

##### Prior marker variance BayesL

The Bayesian Lasso (BayesL) (de Los Campos *et al.*, 2009; Park and Casella, 2008; Legarra *et al.*, 2011) does not set some markers to zero but instead assign them with a very small value that follows the prior distribution:

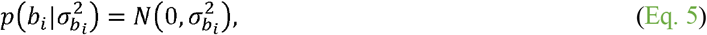

with a Laplace variance distribution,

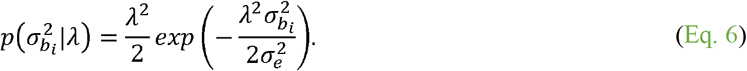

The prior value of *λ* can be expressed as the ratio between marker variance and residual variance (Pérez *et al.*, 2010):

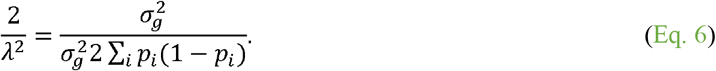

##### Prior marker variance BayesA

Alternatively, one can utilize a priori information, such as the variance of the marker effects 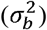. This can be done using a scaled inverted chi-square distribution 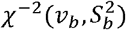, where *S* is a scale parameter and *v* is the number of degrees of freedom (Meuwissen *et al.*, 2001). Using a scale parameter 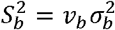, and the prior distribution of marker effects

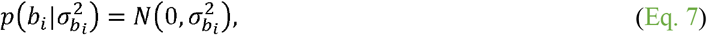

with the prior distribution of variance

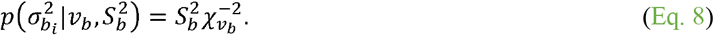

In contrast to BayesN, the prior of marker effects in BayesA follows a t-distribution (Gianola *et al.*, 2009)

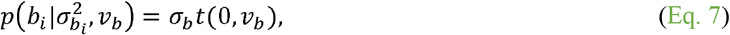

with 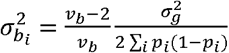, as the variance of a t-distribution is 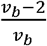.

##### Prior marker variance BayesC

Not all segregating genetic variants in the genome are causal, thus, some individual markers should have zero effect, which is not possible in BayesA as the a priori chi-squared distribution prevents any marker variance from being zero. Therefore, Habier et al. (2011) proposed that some markers have zero effects, while those genetic variants with an effect have a common variance, 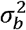. The prior distribution of marker effects is thus,

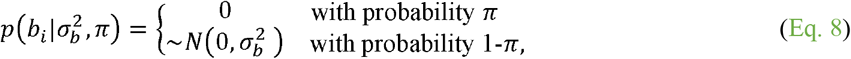

where the prior distribution of the variance is,

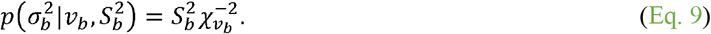

The scale parameter 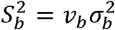, with 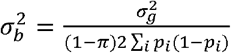. The prior distribution used in BayesC is similar to what is implemented in the commonly used method, LDpred (Privé *et al.*, 2021; Vilhjálmsson *et al.*, 2015).

##### Prior marker variance BayesR

The hierarchical Bayesian model used in BayesC, has been extended to feature additional prior probabilities (Lloyd-Jones *et al.*, 2019; Erbe *et al.*, 2012; Moser *et al.*, 2015). The marker effects, are assumed to be sampled from a mixture with a point mass at zero and univariate normal distributions conditional on marker effect variance 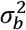 with scaling factors ***γ***:

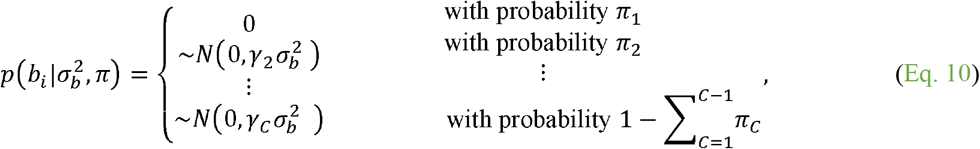

where *π* = (*π*_1_, *π*_2_,…,*τ_C_*) is a vector of prior probabilities and *γ* = (*γ*;_1_, *γ*_2_, *γ_v_*) is a vector of variance scaling factors for each of the *C* marker variance classes. The *γ* scalars are predefined and constrain how the marker effect variance, 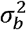, scales within each class distributions. Typically, four classes are used (Lloyd-Jones *et al.*, 2019); *γ* = (0,0.01,0.1,1.0) with the probabilities *π* = (0.95,0.02,0.02,0.01).

##### Estimation of model parameters

Bayesian Linear Regression methods use an iterative algorithm for jointly estimating genetic marker effects. Estimation of the joint marker effects depend on additional model parameters such as the probability of being causal 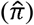, an overall marker variance 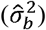, genetic variance 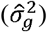 and residual variance 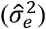, which can be used for estimating trait heritability, 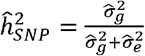. Estimation of the model parameters is obtained using Markov chain Monte Carlo (MCMC) Gibbs sampling from the fully conditional posterior distributions.

#### 3.2.3 Multiple-trait genomic scores

We and others have previously shown, how a genomic predictor can be created as a weighted index combining several GWAS summary statistics, thereby taking advantage of genetic correlations among traits and diseases (Maier *et al.*, 2018; Rohde *et al.*, 2021). In short, the index weights are obtained as:

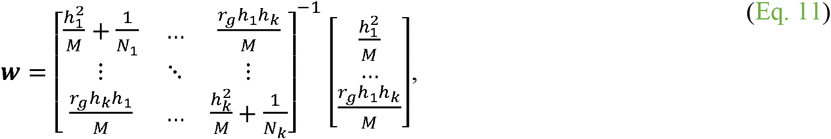

where *h*^2^ and *r_g_* denotes the estimated heritability and genetic correlations, respectively, *M* is the number of independent chromosomal segments (*M* = 60,000 (Yang, Weedon, *et al.*, 2011)), and *N* is the sample size from each individual GWAS. Here, the heritability and genetic correlations were estimated with LDSC as implemented in the qgg package. The multiple-trait genomic score (MT-GS) can then be obtained as the sum of adjusted marker effects 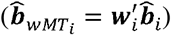:

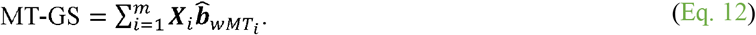

### 3.3 Fine-mapping with BLR models

BLR models used for fine-mapping have been specialized to identify the markers that have the greatest probability of being causal. For BLR models based on mixture normal priors (e.g., BayesC or BayesR), can be used to compute the proportion of iterations from the Gibbs sampling for which a particular marker is included in the model with a non-zero effect, which is referred to as the posterior inclusion probability (PIP) (Schaid *et al.*, 2018). Ranking markers by their PIP is a convenient way to identify putative causal markers. For example, the top *k* markers ranked by their PIP maximize the expected number of causal markers across all possible subsets of size *k.* However, when multiple markers in a region are highly correlated and all are approximately equally associated with the phenotype, it might be better to estimate the posterior expected number of causal markers by summing the estimated PIPs for all markers in a region.

Therefore, for a BLR model that is based on a mixture of normal priors (e.g., BayesC or BayesR), the sum of posterior inclusion probabilities for *m* marker in a window, can be used to make inference on the presence of causal markers in that window. This statistic is referred to as the window posterior inclusion probability (WPIP):

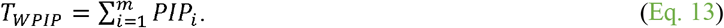

Here, we used sliding window size of 100 markers.

### 3.4 BLR-based gene set enrichment analyses

Aggregating genetic markers into biologically informed entities, such as genes, biological pathways, protein interaction complexes etc., and analysing those sets of markers jointly constitute a valuable addition to single-marker analyses (de Leeuw *et al.*, 2016; Rohde *et al.*, 2016; Sørensen *et al.*, 2017). The qgg package (Rohde *et al.*, 2020) contains a range of different gene set enrichment analyses (GSEA), that when the input data is GWAS summary statistics (in contrast to individual level genetic data) use marginal OLS test statistics or *P*-values. Here, we showcase the utility of using the sum of posterior inclusion probabilities from BayesC in gene sets defined by genes, gene ontologies, pathways, protein complexes, and chemical complexes. The mapping of marker names to genes and gene ontologies (Ashburner *et al.*, 2000) was obtained using the Biocondutor package org.Hs.eg.db (Carlson M, 2019), whereas mapping to biological pathways was done using data from the Reactome database (Gillespie *et al.*, 2022), and mapping to protein complexes and chemical complexes were obtained using data from STRING and STICH, respectively (Szklarczyk *et al.*, 2019, 2016).

Further details on all the statistical genetic models presented through the **Methods**, can be found at our accompanied homepage: www.qganalytics.com.

## 4 Results

Here, we present a major update of the R package qgg (Rohde *et al.*, 2020), that provides a unified framework for efficient processing of large-scale biobank genetic data and user-friendly interface for quantitative genetic analysis of complex traits and diseases.

### 4.1 Genomic prediction of standing height and body mass index

Utilising phenotypic and genetic data from the UKB (Bycroft *et al.*, 2018) the white-British unrelated sub-cohort was split into five random training cohorts each containing 300,000 individuals. First OLS association tests between trait phenotype and quality-controlled genotype variants were performed for each sex separately, and combined, within each training cohort.

The genetic marker effects 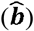 from each training cohort was used to construct sex-specific genomic scores (GS) for the individuals not included within the training cohorts. Expectedly, the *P*-value thresholds used in clumping and thresholding (C+T) showed different prediction accuracy optima for the two traits (Figure 2), and both BayesC and BayesR outperformed the other Bayesian shrinkage models and C+T (Figure 2). No difference in predictive performance between sexes was seen for BMI (Supplementary Fig. 1), however, for standing height, the prediction accuracy was significant higher for females compared with males when using the BLR models (Supplementary Fig. 1).

**Figure 2.**
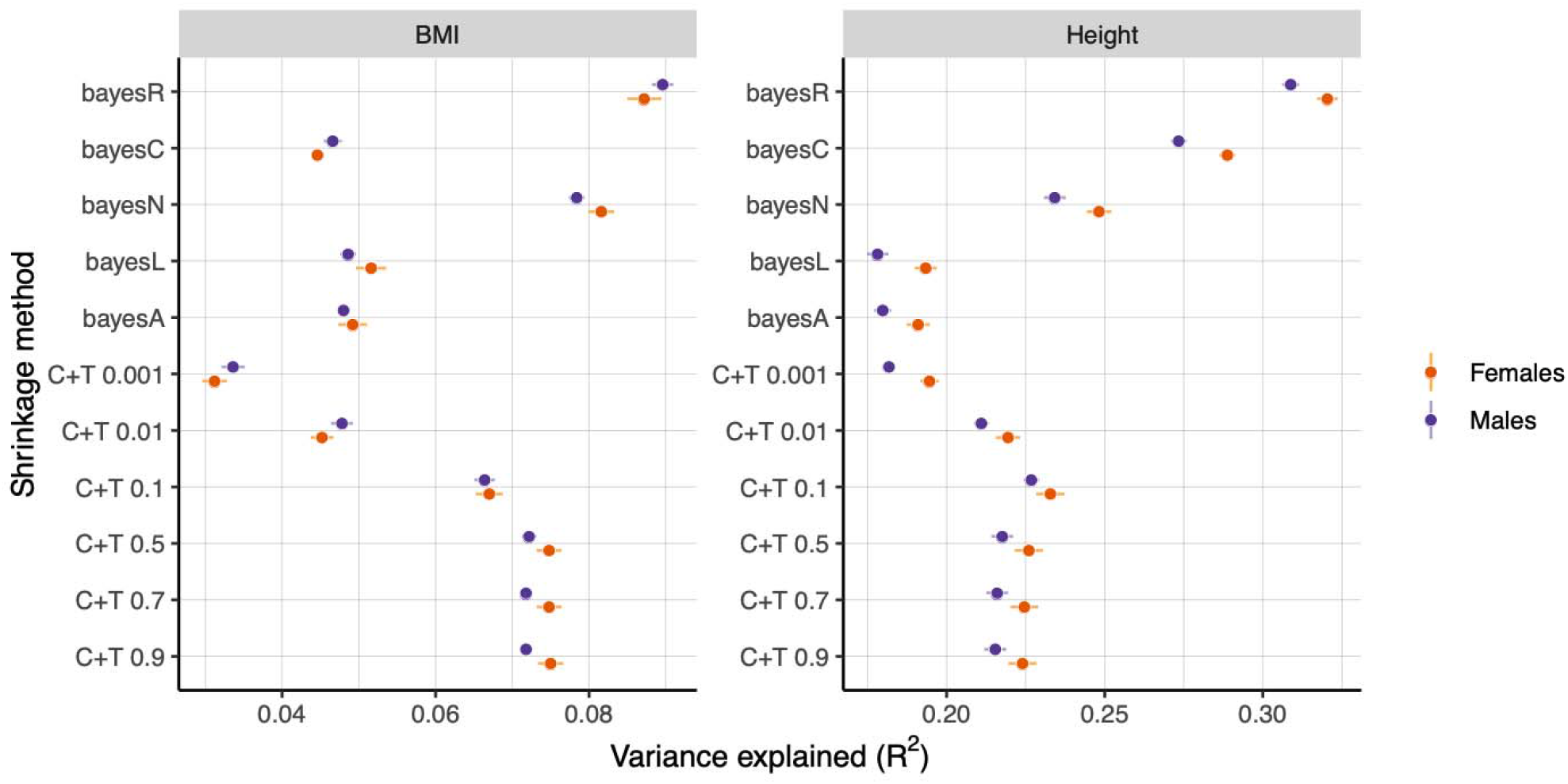
Average prediction accuracies (error bars are standard errors of the mean) for body mass index (BMI) and standing height for males and females separately quantified as variance explained (R^2^). Different shrinkage algorithms were tested including clumping and thresholding (C+T,), and five different Bayesian Linear Regression (BLR) models.

Next, we computed multiple-trait GS by first constructing index selection weights (Eq. 11). The multiple-trait GS increased the prediction accuracy for BMI for males and females with >20% (Figure 3), despite the low genetic correlation between height and BMI (Figure 3). Similarly, leveraging the genomic pleiotropy between BMI and height for males and females, separately increased the prediction accuracy for standing height by almost 15% (Figure 4).

**Figure 3.**
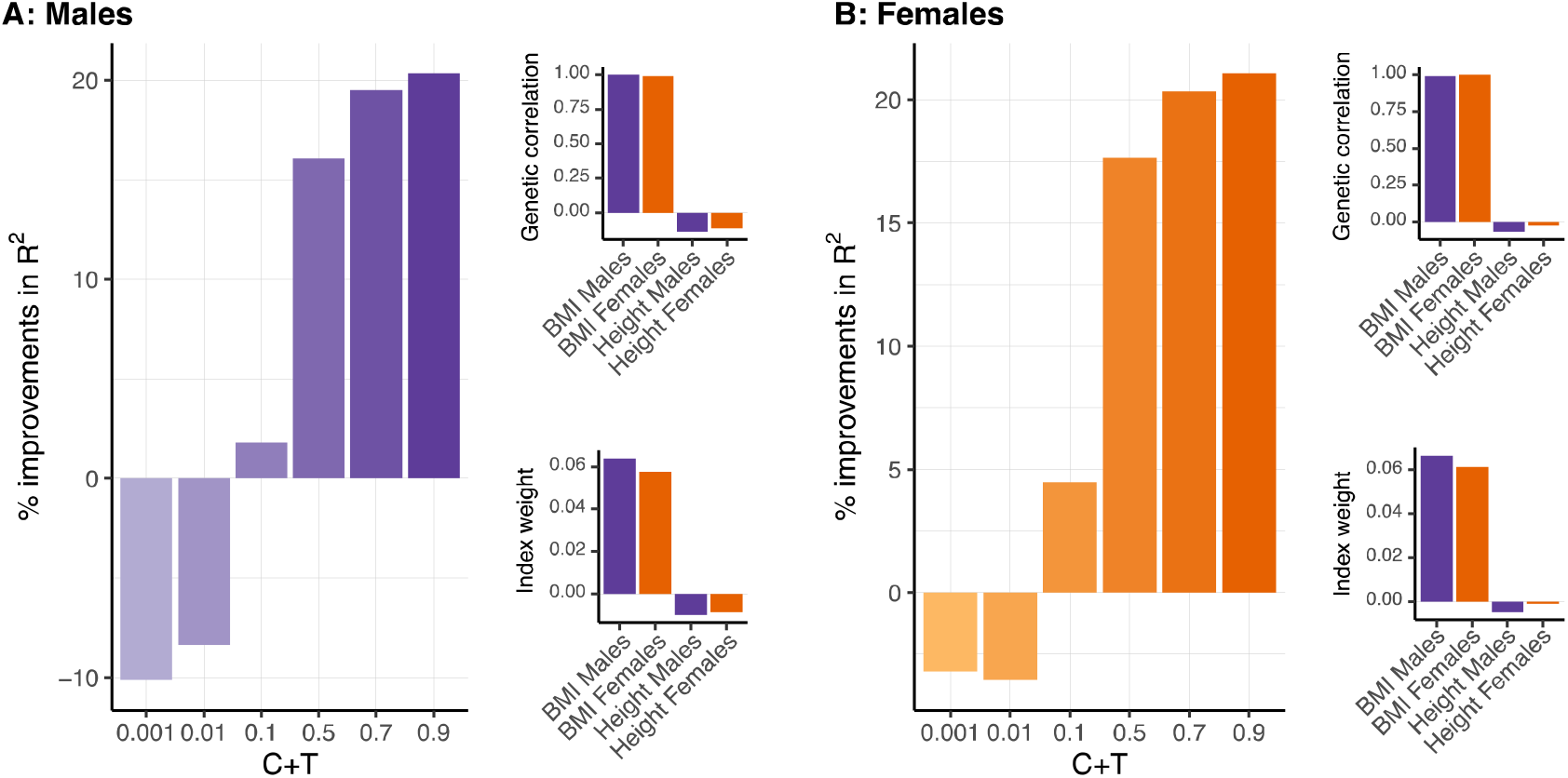
Improvements in predictive performance for body mass index (BMI) for **A**: males and **B**: females, when leveraging the genetic correlation between BMI and standing height for males and females separately.

**Figure 4.**
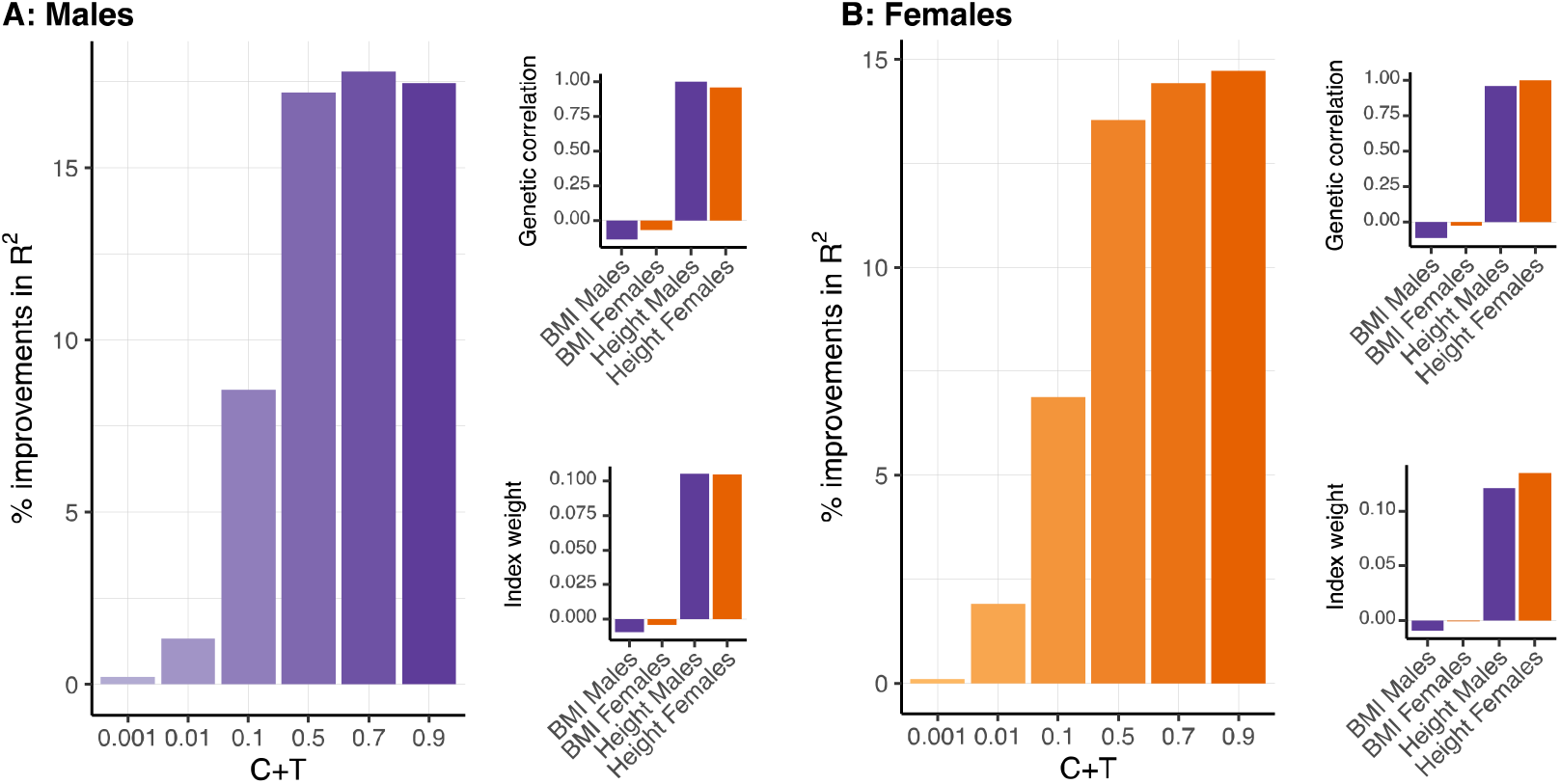
Improvements in predictive performance for standing height for **A**: males and **B**: females, when leveraging the genetic correlation (estimated with LDSC) between BMI and standing height for males and females separately.

Applying the selection index MT methodology to the BLR models further improved the predictive accuracy for all five BLR models with 20-60% for body mass index and 10-45% for standing height (Figure 5A). Interestingly, MT-GS based on BayesR displayed same amount of prediction accuracy as single trait GS with BayesR when the summary statistics are trained on both males and females (Figure 5B), i.e., approximately double the sample size to the analysis of males and females separately.

**Figure 5.**
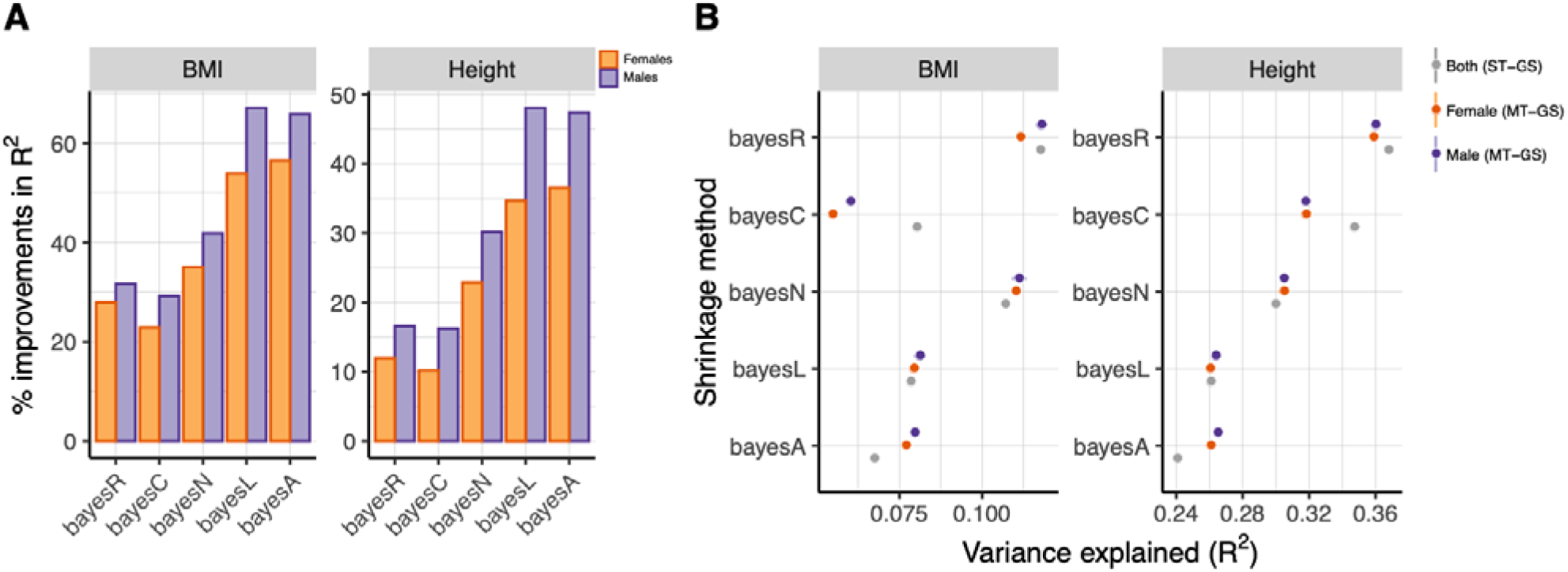
Average prediction accuracies of the five different Bayesian Linear Regression (BLR) models (error bars are standard errors of the mean) for body mass index (BMI) and standing height for **A**) males and females separately using the multiple-trait approach (MT-GS), and for **B**) both sexes combined using the single-trait approach (ST-GS) quantified as variance explained (R^2^).

### 4.2 Estimated genomic parameters

Computing the index weights (Eq. 11) used in constructing the multiple-trait genomic scores (Eq. 12) requires estimates of trait heritability () and the genetic correlation () among the phenotypes, which can e.g., be achieved using summary statistics-based LD score regression (as implemented in qgg). However, our implementation of the different BLR models also provide accurate estimates of the heritability. For example, using BayesC we estimated the heritability for BMI and standing height for each sex separately and the two sexes combined. The heritability for height and BMI were within trait the same across sexes and combined with very small differences in estimates across the five training sets (Figure 6A).

**Figure 6.**
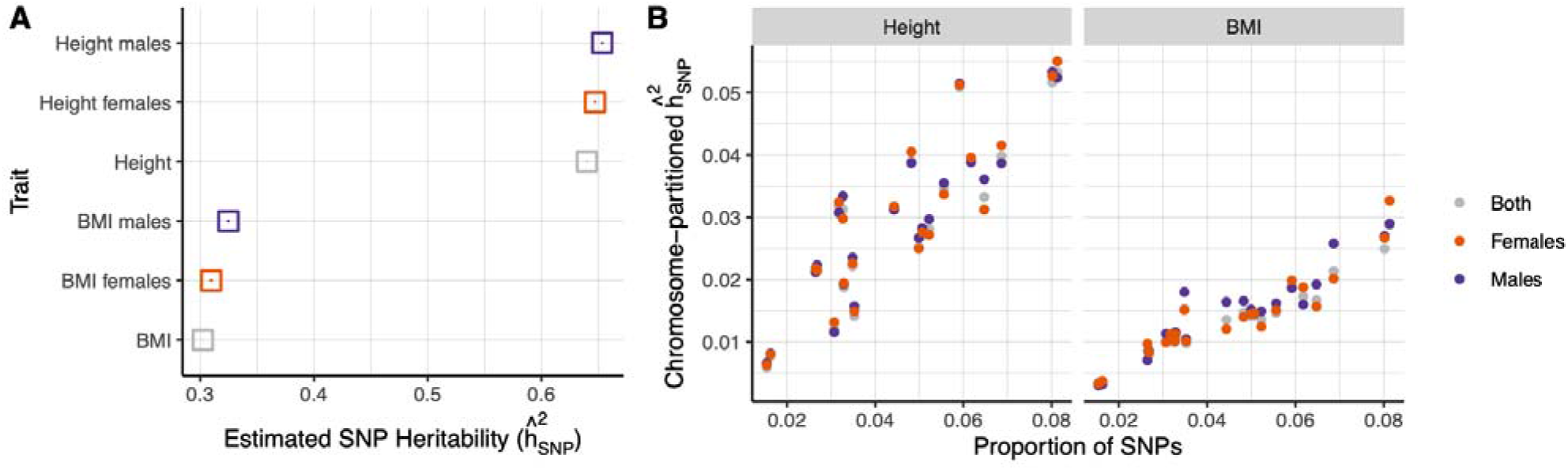
Bayesian linear regression (BLR) model with bayes R shrinkage for **A**) SNP heritability estimation () for standing height and body mass index (BMI) for each sex separately and together, and **B**) partitioned heritability into autosomal chromosomes.

Furthermore, we can also partition the heritability across different marker sets, e.g., by those on each autosomal chromosomes (Figure 6B). In addition, the genetic correlation can also be partitioned, which showed varying degree of positive genetic correlation across the 22 autosomes for standing height between sexes (Figure 7). For BMI the pattern was different with almost the same level of genetic correlation among chromosomes (Figure 7), and the genetic correlation between BMI and height between sexes showed different patterns across autosomes.

**Figure 7.**
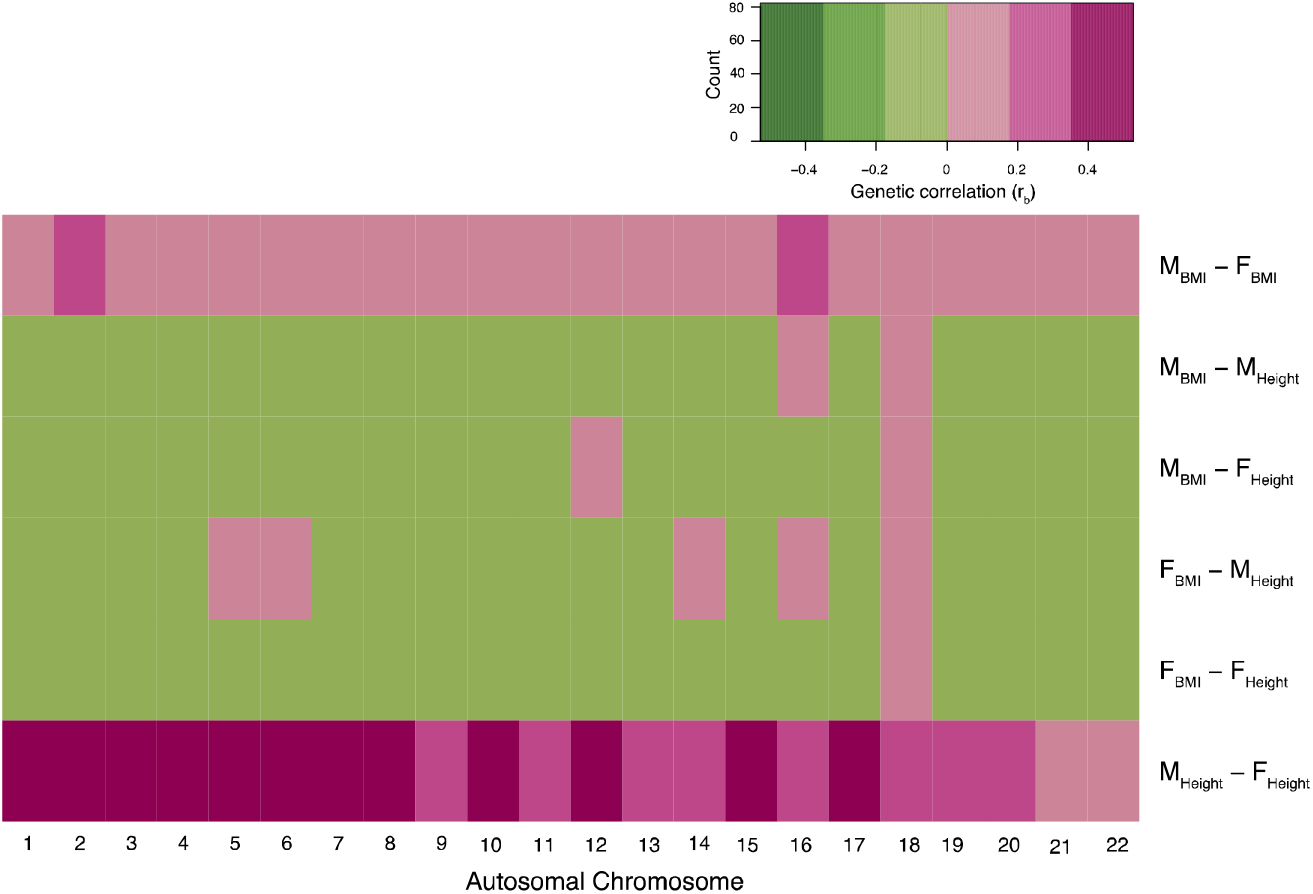
Partitioned genetic correlation (here approximated as the correlation among BayesC estimated marker effects, among autosomal chromosomes for standing height and body mass index (BMI) for males and females.

Besides estimating how much of the phenotypic variance we can explain with common genetic variants, or how large a fraction of the genetic contribution that is shared across traits, it can also be useful to estimate the proportion of variants with a likely causal effect. As example, using BayesR, we estimated the proportion of variants with no, small, moderate, or large effect for the two traits. BMI and height, has to a large extent a similar genetic profile with one big difference, namely that BMI have <5% variants with large effects, while standing height has ~15% variants with an estimated large effect (Figure 8).

**Figure 8.**
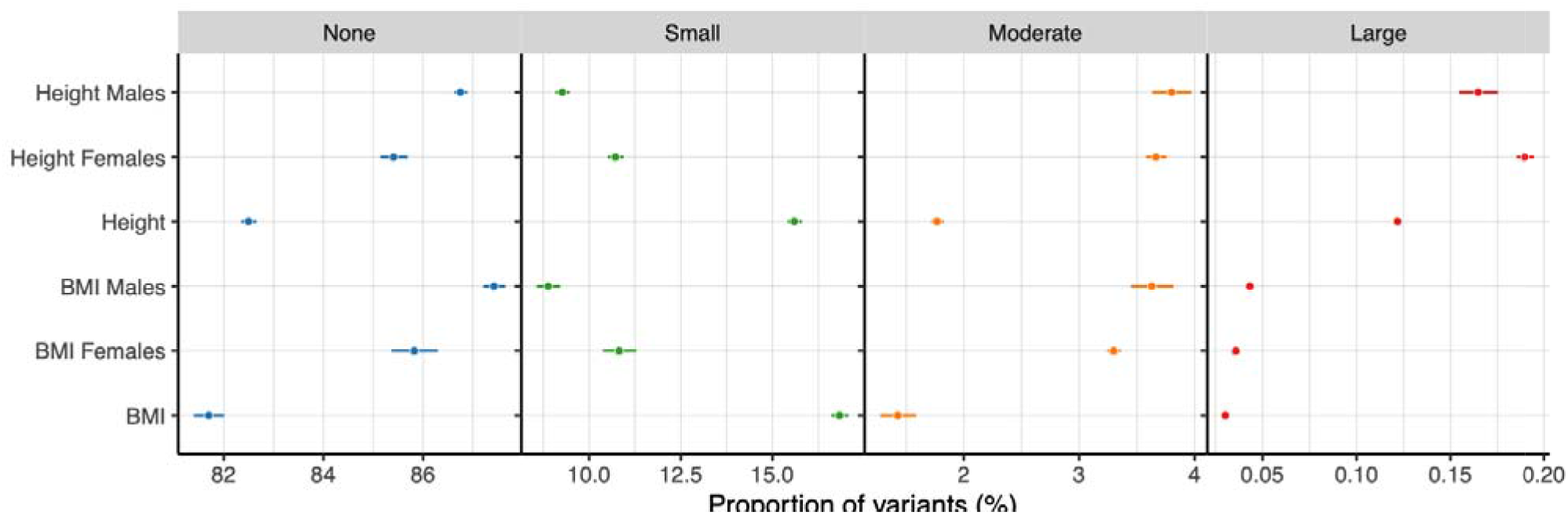
Proportion of genetic variants with no, small, moderate, or large effect sizes, estimated with BayesR. Each point is the mean across the five training sets, and error bars represents the standard error of the mean.

### 4.3 Bayesian fine-mapping

BLR models account for the underlying genetic architecture by sampling marker variances from different prior distributions and by accounting for linkage disequilibrium between markers these models are proposed to have greater power to detect causal associations (Lloyd-Jones *et al.*, 2019). As example, we used BayesC to fine-map genetic markers for standing height, focusing on chromosome 1. Expectedly, because of the BayesC prior shrinkage, most of the markers on chromosome 1 received a zero-effect size (Figure 9A). We computed the window posterior inclusion probability (WPIP) for a sliding window of 100 markers to make inference on the presence of causal markers in these windows. Particular one region on chromosome 1 showed high WPIP compared to the other regions (Figure 9B). This region contains the *PAPPA2* gene which is one of the many known QTLs for standing height (Figure 9C) (Allen *et al.*, 2010; Kichaev *et al.*, 2019; He *et al.*, 2015). Likewise, zooming in on another region with lower WPIP than the *PAPPA2*-region, we find other regions with high posterior inclusion probabilities, such as the intergenic region between the genes *H6PD* and *SPSB1,* which also are known genes for height (Berndt *et al.*, 2013; Sakaue *et al.*, 2021; Wood *et al.*, 2014).

**Figure 9.**
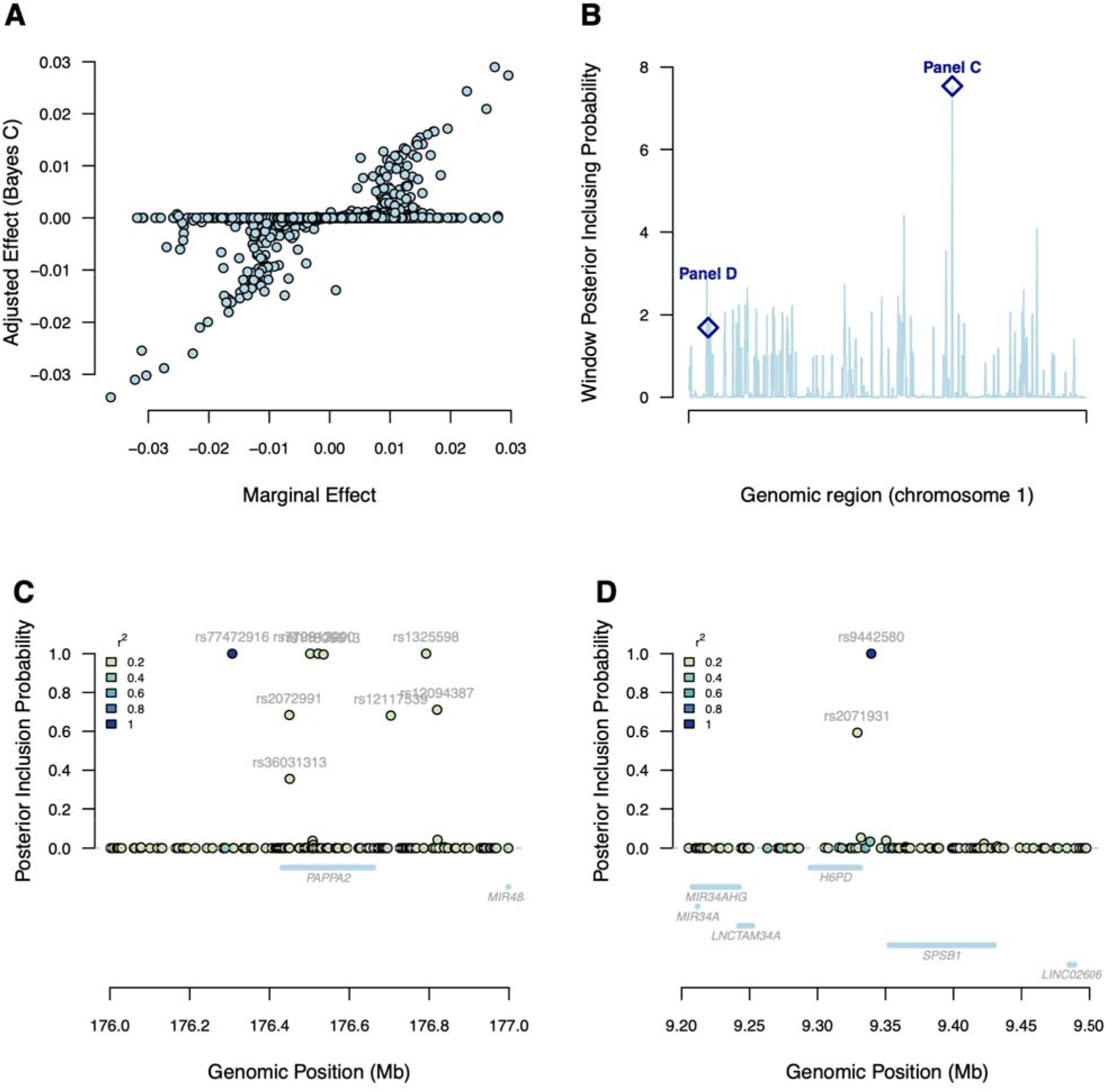
Example of genetic fine mapping using a Bayesian linear regression (BLR) model with Bayes C shrinkage for standing height on chromosome 1. **A**) Comparison of ordinary linear squares (OLS) marginal SNP effects to adjusted SNP-effects after Bayes C adjustment. **B**) Chromosome 1 Manhattan-like plot of windows of posterior inclusion probabilities (Bayes C, sliding window size of 100 SNPs) for standing height. **C-D**) Locus zoom on two genomic regions on chromosome 1 (see diamond indicators in panel B) of individual variants’ posterior inclusion probability. Color grading indicates degree of linkage disequilibrium (LD) measured by Pearson’s correlation (r^2^). Blue segments below points indicate gene transcription start and stop sites (UCSC, GRCh73).

### 4.4 Bayesian-based gene set enrichment analysis

An overview of the number of identified genetic marker sets with a false discovery rate below 5% is shown in Table 1. A few examples of association are given below, while a complete overview can be found in Supplementary Table S1 and at our accompanied homepage (qganalytics.com). We identified *ZNF385C* and *CDIP1* as being associated with standing height for both males and females, while *ANKS6* was female-specific for height. For both sexes and traits, we show an enrichment in the sequence ontology ‘coding region’. For gene ontologies we identified a total of 13 terms that were associated with height in both sexes, including terms within the molecular function for ‘negative regulation of transcription by RNA polymerase II’ (GO:0000122), ‘positive regulation of DNA-templated transcription’ (GO:0045893), and ‘DNA-binding transcription factor activity, RNA polymerase II-specific’ (GO:0000981). Only a few Reactome pathways were identified, in particular DNA synthesis (R-HSA-164516 and R-HSA-164525) for BMI for males. None of the chemical complexes survived correction of multiple testing.

**Table 1.** Summary of number of marker sets within the different categories with an FDR corrected *P*-value <0.05.

| Gene set (total sets) | Height |  | BMI |  |
| --- | --- | --- | --- | --- |
|  | Males | Females | Males | Females |
| Sequence ontology (7) | 5 | 5 | 0 | 1 |
| Genes (24,059) | 2 | 3 | 5 | 1 |
| Gene ontologies (4,544) | 13 | 13 | 0 | 2 |
| Pathways (2,243) | 0 | 1 | 4 | 0 |
| Protein complexes (4,948) | 0 | 1 | 1 | 0 |
| Chemical complexes (94,884) | 0 | 0 | 0 | 0 |

## 5 Discussion

Here, we have presented a major update of the R package qgg for quantitative genetic and genomic analyses of complex traits and diseases. Particularly, we focused on the expanded utilities relating to genomic prediction using Bayesian linear regression models. It has repeatedly been argued that because BLR models fit all genetic markers simultaneously while accounting for LD among genetic variants, and because marker variances are sampled from pre-specified prior distributions, these models have greater statistical power to identify causal genetic variants (Moser *et al.*, 2015; Lloyd-Jones *et al.*, 2019; Vilhjálmsson *et al.*, 2015; Ehsani *et al.*, 2012, 2016; Sørensen *et al.*, 2015), entailing increased prediction accuracy. We observed for both body mass index and standing height, that BLR using bayesR was superior to the compared models (Figure 2), however, the ranking of the other methods was trait specific (Figure 2), supporting the notion that different shrinkage methods perform differently on different traits most likely because of different underlying genetic architectures.

Our reported prediction accuracies were comparable to what was reported by Lloyd-Jones et al. (2019) whom first presented the bayesR methodology using GWAS summary statistics. Other commonly used tools for constructing whole-genome risk scores, such as LDpred (Privé *et al.*, 2021; Vilhjálmsson *et al.*, 2015) and PRsice (Euesden *et al.*, 2015), has despite their great popularity limited applications as they rely on a single analytical approach, namely that LDpred utilise bayesC methodology and PRsice is based on C+T. As the genetic architecture of complex traits differs for different traits and diseases, it is most likely that there do not exist a single best algorithm for constructing genomic predictors. Therefore, having statistical tools that offers different predictors might be valuable; exactly what qgg offers.

Generating multivariate genomic predictors derived from multiple GWAS summary statistics of correlated phenotypes increase prediction accuracy (Maier *et al.*, 2018; Rohde *et al.*, 2021), and here we illustrate that combining bayesR with the weighted index of correlated traits have the same predictive ability as a predictor based on single GWAS summary statistics with a sample size the double of the multiple-trait predictor (Figure 5B). This clearly demonstrate the added value of combining several, potentially smaller-sized GWAS’es, to generate accurate genomic predictors.

A further added value by our implementation of the BLR models is accurate estimation of genetic parameters, such as heritability (Figure 6), and increased power to fine map causal genetic variants (Figure 7), which remain absent in many other commonly used tools for genetic risk prediction. Also, we provide examples on how BLR models can be used to fine map putative causal variants. The challenge of BLR models is determining which markers have non-zero effect sizes on a trait. The BLR models are fitted using all markers simultaneously, and fine mapping approaches must therefore consider sets of markers in a genomic region rather than individual markers, because the effect of a causative mutation may be distributed across multiple markers. Although markers with non-zero effect sizes may be referred to as causal, it is important to realize that statistical methods alone cannot determine causality.

Therefore, we used a statistic we refer to as window posterior inclusion probability to identify genomic regions associated with the trait phenotype. Simulation studies has shown that statistics such as WPIP are superior to other methods in terms of power to detect causal loci for both traits with oligogenic and polygenic architectures (Lima *et al.*, 2022). Furthermore, fitting all markers in the region results in a concentration of the association signals to a small region around the causal loci with control of the posterior type I error rate (Fernando *et al.*, 2017). For BLR models, several association statistics have been developed to identify which markers or windows of markers can be considered as explaining a substantial or significant proportion of the genetic variance. However, further studies are required to determine optimal association statistics for the BLR models.

Finally, multiple trait BLR models can be used to better understand and for dissecting genetic correlations between traits using genetic markers, e.g., evaluating whether pleiotropy or linkage disequilibrium are at the root of between trait associations. In the new release of qgg we have implemented a general multiple trait BLR model based on the BayesC prior (Cheng *et al.*, 2018). This model is particularly interesting because it provides insight into whether markers affect all, some, or none of the included traits. However, what the impact is of the choice of prior assumptions on marker effects across traits in currently unknown and requires further investigation. Also, an important limitation of all methods that attempt to use genotype–phenotype associations to detect pleiotropic loci is that they cannot differentiate the presence of a pleiotropic locus from the presence of two closely linked single-trait loci, depending on the extent of LD in the region. Furthermore, issues with inferences about genetic (co) variance based on BLR models can misrepresent the true genetic parameters if the causal loci are not genotyped because of incomplete LD between markers and causal loci and among causal loci (Gianola *et al.*, 2015; de los Campos *et al.*, 2015).

## 6 Conclusion

Here, we have shown a multitude of the new functionalities implemented in the qgg package. In particular, we highlight the five BLR models for single trait and multiple trait analyses for 1) constructing genomic scores with utilities in genomic medicine, 2) fine mapping of GWAS results, and 3) accurate estimation of quantitative genetic parameters.

## 7 Software and code availability

The methods presented in the study is implemented in the R-package qgg and is available from CRAN (https://cran.r-project.org/web/packages/qgg/index.html), or as a developer version at our GitHub page (https://github.com/psoerensen/qgg). Detailed notes on the implemented statistical genetic models, tutorials and example scripts are available from our accompanied homepage https://qganalytics.com/.

## Acknowledgements

The data used in the presented study were obtained from the UK Biobank Resource (project id 31269). All of the computing for this project was performed on the GenomeDK HPC cluster. We would like to thank GenomeDK and Aarhus University for providing computational resources and support that contributed to these research results.

