## Supplementary Material for "Expanded utility of the R package qgg with applications within genomic medicine"

^2^Centre for Quantitative Genetics and Genomics, Aarhus University, Aarhus, Denmark
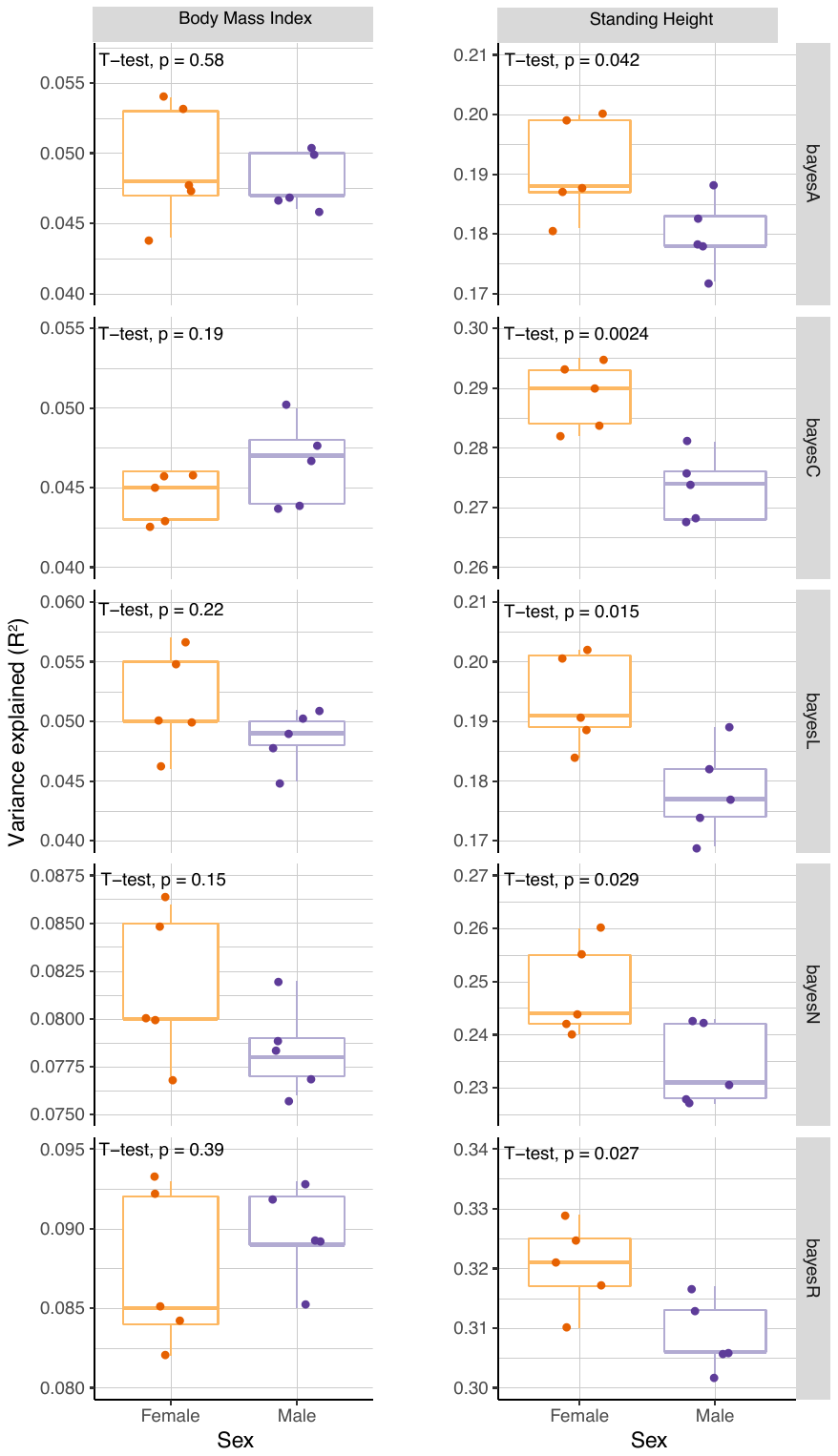

**Supplementary Figure S1.** Boxplot of predictive performances (quantified as variance explained, R^2^) for body mass index and standing height within the UK Biobank across the five training/validation cohorts. The T-test P-values is testing differences in mean predictive performance between females and males.
